# Unbiased emission factor estimators for large scale forest inventories: domain assessment techniques

**DOI:** 10.1101/138131

**Authors:** Luca Birigazzi, Javier G. P. Gamarra, Timothy G. Gregoire

## Abstract

Large scale forest inventories are often undertaken following a stratified random or systematic design. Yet the strata rarely correspond to the reporting areas of interest (domains) over which the country wants to report specific variables. The process is exemplified by a country aiming to use national forest inventory data to obtain average biomass estimates per forest type for GHGI international reporting, where activity data (areas of land use or land use changes) and emission factors (carbon coefficients) are typically compiled from disparate sources and estimated using different sampling schemes. This study aims to provide a decision tree for the use of data obtained from forest surveys to draw conclusions about population sub-groups created after (and independently of) the sample selection. While bias can arise whenever activity data and emission factors are calculated independently, it can be eliminated in case of a simple random or simple systematic design if properly weighted estimators are provided. This manuscript describes two unbiased estimators that can be used to estimate reporting-strata means, regardless of the sampling design adopted, and extends the result to the common situation in which the reporting-strata are spatially explicit, where a nested group estimator outperforms in terms of both bias and precision other more traditional estimators. From this estimator, an optimal sample allocation scheme is also derived.

## 1 Introduction

Country-specific estimates of carbon coefficients (aka *emission factors*) are required to compile national greenhouse gas inventories (GHGI) under the United Nations Framework Convention on Climate Change (UNFCCC) and, in the context of REDD+, for developing Forest Reference Levels (FRLs) and for the reporting of REDD+ results-based actions. To this end, in order to account for emissions from land use, land use change and forestry (LULUCF), different emission factors have to be estimated for each of a number of land use categories and of various other land subpopulations, typically defined according to climatic zone, forest type or management practices (cf. IPCC 2003, 2006). Notice that these estimates are required to be unbiased and as precise as possible^1^.

IPCC tier 3 methods, which, if well implemented, are supposed to be the most certain and reliable, require these forest carbon coefficients to be obtained from national forest inventories (NFI). Due to the intrinsic variability of biomes and land uses in most of the countries, sampling designs for NFIs tend to rely on stratification as a first step to reduce uncertainties in emission estimates (Köhl et al 2006; Maniatis and Mollicone 2010). But the subpopulations for which the emission factors are needed might differ from those designated at the planning phase of the forest inventory. That is, the requested reporting units or areas of interest for the emission factors might be identified after the definition of the sampling design and/or after the data collection has been carried out. These targeted subpopulations are often called *domains of interest* in the statistical literature (Cochran 1977; Särndal et al 1992; Särndal and Lundström 2005; Köhl et al 2006; Gregoire and Valentine 2008; Mandallaz 2008; Lohr 2009; Schulz et al 2009; Thompson 2012). The lack of congruency between strata and domains usually results in a random and often small number of observations for each emission factor. The situation can be made even more complicated in the presence of complex forest inventory survey designs, often not optimized for all variables of interest involved in multipurpose inventories. In all these cases specific statistical approaches must be adopted in order to ensure that the estimates are precise and unbiased, very often requiring ancillary data or model-, rather than design-based inference (Schreuder et al 2004; Chanbers 2011). Similar issues might arise whenever estimates of forest carbon are required not only at the national level, but also for certain provinces or districts of the country (Rao and Molina 2015).

Salient features of the estimation of emission factors for the LULUCF sector are that the domains are often defined spatially over a landscape and that their sizes are usually known but frequently obtained independently of the NFI. The emission factors, in this case, would consist of the average values of carbon or biomass for those specific areas. A detailed review of basic estimation methods for domains is provided in §10.3 of Särndal et al (1992). A compilation of domain estimators used for the analysis of the U.S.A. forest inventory data is presented in Bechtold and Patterson (2005) and for the Swiss National forest inventory by Mandallaz (2008).

Here we aim to provide an overall vision of the different domain estimation techniques applicable to derive LULUCF emission factors from large scale forest inventories. The focus here is particularly on NFI data which are often available to/in REDD+ countries.

Our results apply both to cases in which domain sizes are estimated within or independently of the NFI. In the case where the domains are known and spatially explicit we also derive an unbiased estimator that improves the standard errors typically associated to classical π-weighted (i.e., Horwitz-Thompson) estimators, as well as the optimum sampling sizes per stratum derived from this estimator. Finally, we propose a decision tree, targeted particularly to countries aiming to report emission factor estimates from national forest inventories, to select the most appropriate domain mean estimator.

## 2 General formulation for domain estimators

Let the population of interest be denoted by *𝒫*, and the size of *𝒫*, as measured by the number of areal units of uniform size, be denoted by *N*, so that *𝒫* = {*u_1_*,…, *u_k_*,…, *u_N_* }. Let *𝒫* be partitioned into *H* non-overlapping subpopulations *𝒫_1_*,…, *𝒫_H_*. These will be referred to as the sampling strata. Let *N_h_* denote the size of *𝒫_h_*, with *h = 1,…,H* such that 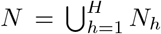. The population may also be partitioned alternately into *D reporting strata*, known widely in literature on survey sampling as *domains*, cf. Särndal et al (1992, Chap. 10). To this end, let *𝒰_d_* denote the *d^th^* domain. Let *N_d_* denote the size of *𝒰_d_, d* = 1,…, *D* such that 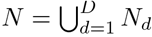 and let *𝒫_d_ = N_d_/N* denote the relative size of the domain *𝒰_d_* (Table 1). In this paper we address the estimation of the domain averages, *Y_d_*, for some variable on interest indicated by *Y*. Implicit in the above is that each population unit can be a member of a single sampling stratum and domain.

**Table 1.** definitions of weights and proportions

| Symbol | Definition |
| --- | --- |
| $P_d$ | $N_d/N$ |
| $Q_d$ | $1 - P_d$ |
| $W_h$ | $N_h/N$ |
| $P_{hd}$ | $N_{hd}/N_h$ |
| $Q_{hd}$ | $1 - P_{hd}$ |
| $W_{hd}$ | $N_{hd}/N_d$ |
| $p_{hd}$ | $n_{hd}/n_h$ |
| $f_d$ | $n_d/N_d$ |
| $f_h$ | $n_h/N_h$ |

Let *s_d_* define the part of the sample that happens to fall into *𝒰_d_*, so that: 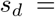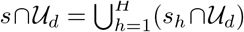. Using results that are elaborated in §5.8, 7.6, and 10.3 of Särndal et al (1992) (see also Cochran 1977; Mandallaz 2008; Thompson 2012), an approximately design-unbiased estimator for the domain mean, whether or not the domain size is known, is

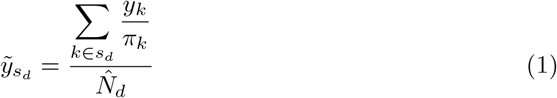

where *π_k_* is the inclusion probability of the *k^th^* unit *u_k_*, and 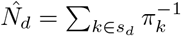. The approximate variance of 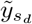 is given by

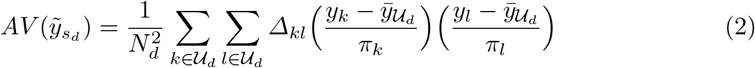

where 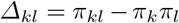 is the covariance between the *sample membership indicators*. An estimator of *AV*(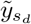) is

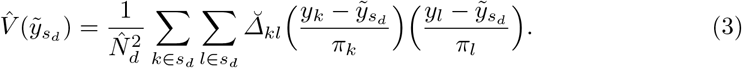

where 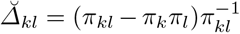. Specific applications of these general formulas to the most common sampling designs are provided in the next sections. The underlying assumption is that the probability that *s_d_* is empty is negligible.

In the following sections we will assess specific applications of these general formulas to the most common sampling designs: simple random and stratified random sampling.

## 3 Simple Random Sampling

Let us consider the case in which a simple random sampling (SRS) is carried out on the population *𝒫*. That is, a sample *s* of size *n* is randomly selected from *𝒫*. Let us define *n_d_* the size of the sample falling in the domain *𝒰_d_*. Under simple random sampling (where *π_k_* = *n/N*), given the event *n_d_* > 0, the estimator 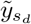 (Eq. (1)), is approximately design-unbiased for the domain mean and corresponds to the domain sample mean, namely

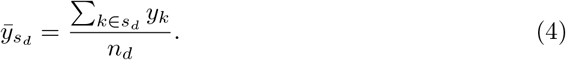

However, when the probability that *n_d_ = 0* is not negligible, the bias of 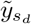 may be substantial. The same result applies also to the case in which a systematic sampling is carried out (with *π_k_* = 1/*a*, where *a* is the sampling interval between successively sampled units). From Eq. (2), the approximate variance of 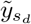 is

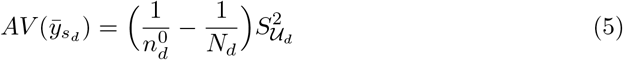

where 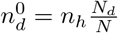 is the expected domain sample size and 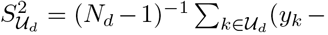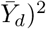 is the domain variance. Eq. (5), however, is likely to underestimate the variance of 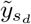, due the fact that, under a simple random sampling, the domain sample size *n_d_* is a random variable, such that 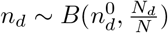. Each time a sample of size *n* is drawn from the population, the number of sample units that fall in the domain *𝒰_d_* might differ (or even be zero). The random domain sample size results in a loss of precision, which is inversely proportional to sample size n and to the domain proportion *P_d_.* Conditioning on *n_d_* > 0 provides a better approximation of the variance of 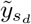, namely

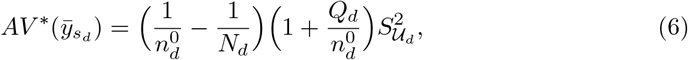

where *Q_d_* = 1 — *P_d_*, considering binomial probabilities, as shown in §10.4 of Särndal et al (1992). Notice that Eq. (6) requires the domain size to be known.

The variance estimator in Eq. (3) becomes

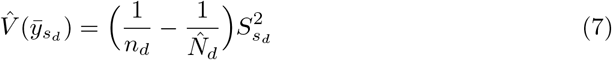

where 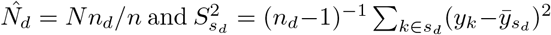 is the domain sample variance.

## 4 Stratified Random Sampling

Let us consider the case in which a stratified random sampling is carried out on the population *𝒫*, based on the *H* sampling-strata. That is, a probability sample *s_h_*, of size *n_h_*, is independently selected from each *𝒫_h_*, according to a design *p_h_*(). The total sample set, denoted as *s*, will be composed as: 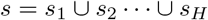.

Any stratum will intersect a certain number of domains. Let us define the intersection among the *d^th^* domain and the *h^th^* sampling-stratum as 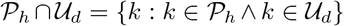 and let *N_hd_* denote the size of 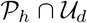 We can now define the following proportions: *W_h_ = N_h_/N*, which is the stratum weight, *W_hd_ = N_hd_/N_d_*, which is the intersection weight within the domain, and *P_hd_ = N_hd_/N_h_*, which is the relative size of the intersection within the stratum. The definitions of weights and proportion used in this paper are displayed in Table 1.

### 4.1 Bias of 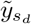 under stratified designs

The arithmetic mean of *Y* for *s_d_* (Eq. (4)) is certainly one of the simplest estimator of the domain mean, but it may be significantly biased under stratified random or stratified systematic designs (Pacificador Jr 1997). A notable exception is constituted by stratified designs with sample allocation proportional to the size of the strata (Holt and Smith 1979). In this case the estimator 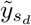 (Eq. (1)) corresponds to the arithmetic sample mean 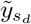 (Eq. (4)), as long as n >> 0, to avoid spurious rounding off effects due to limited allocated sample size. In general, under stratified designs, the magnitude of the bias of 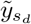 will depend on the allocation of the sample units among the strata and on the homogeneity of the domains (Kish 1980). The development for the quantification of bias in domains under stratified random designs is provided in Appendix A).

### 4.2 Estimators under unknown domain size

Let denote *s_hd_* the intersection of the domain *𝒰_d_* and the sample drawn in the stratum *𝒫_h_*, and *n_hd_* denote the size of this subsample. Under a stratified random sampling, with *H* strata intersecting *D* domains, Eq. (1) can be written as:

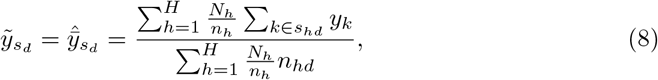

as shown in Särndal et al (1992, example 10.3.3).

Let us call 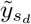 in Eq. (8) π-estimator. It might be informative to re-define the right-hand side of Eq. (8) as

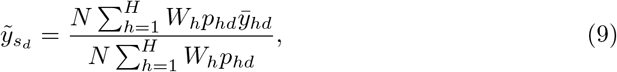

where 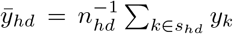 is the arithmetic mean in *s_hd_*. This reformulation makes more explicit that 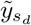 is the ratio of an estimator of the population total of *y* in *𝒰_d_* to an estimator of *N_d_*.

An estimator of the variance of 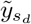 is

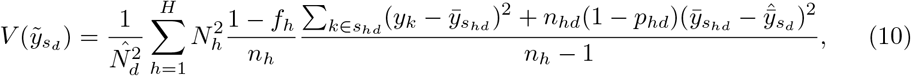

where 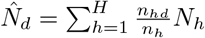, as in Särndal et al (1992, example 10.3.3).

### 4.3 Estimators under known domain size

#### Spatially explicit domains: a nested group estimator

In situations in which it is possible to classify each population unit into a domain, so that the size *N_hd_* of each intersection is known, we propose an alternative estimator of domain means. In these cases, an unbiased estimator of the mean of the domain *𝒰_d_* can be obtained considering the domain *𝒰_d_* itself divided into further strata, given by its intersections with the sampling-strata, (which are 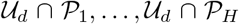), so that 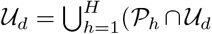, and subsequently applying the usual stratified sampling estimator. The weight of each “nested stratum” is given by the ratio of the size of the intersection to the total size of the domain. This estimator for the mean of the domain *𝒰_d_* is 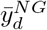(*NG for nested group*):

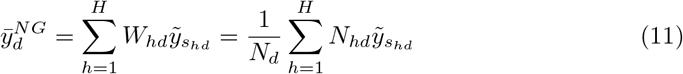

where*W_hd_*=*N_hd_/N_d_* and

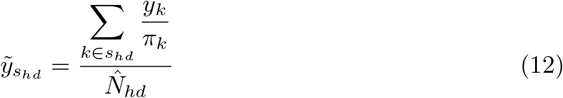

is the π-weighted intersection sample mean, with 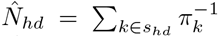. Eq. (11) is a special case of the post-stratified domain estimator (cf. Särndal et al 1992, Eq. 10.7.4 and Remark 10.7.3) for stratified designs, in which the strata are used as post-strata.

If a random or systematic sample has been selected in each stratum, the estimator 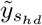 in Eq. (12) is identical to the arithmetic sample mean 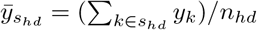, and Eq. (11) becomes

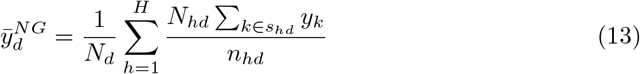

It is worthwhile to notice that 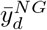 requires *n_hd_* > 0, while the n-estimator - Eq. (9) -requires only *n_h_ > 0*.

As in the previous development in §4.2, it is informative to re-define the right-hand side of Eq. (13), which simplifies further into:

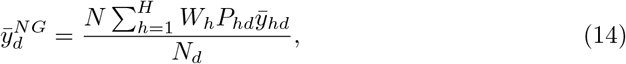

because it makes more evident that, unlike Eq. (9), 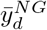 is not a ratio of two estimators, but rather a ratio of an estimator of the domain total to the actual size of the domain. Notice also that the sample estimator *p_hd_* in the numerator of (9) is substituted here by the known value of the relative size *P_hd_*.

The variance of 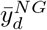 is

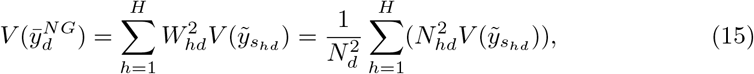

where *V*(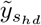) is the variance of the estimator 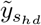.

If a random sample has been selected in each stratum, *V* (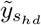) can be re-expressed by analogy to Eq. (6) as

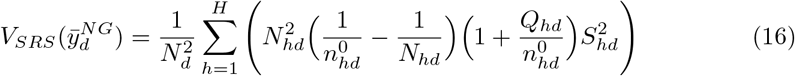

where 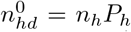 is the expected domain intersection sample size, *Q_hd_ =1 — P_h_* and 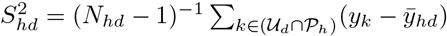 is the intersection variance.

By analogy to Eq. (7), an estimator of *V_SRS_*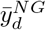 is

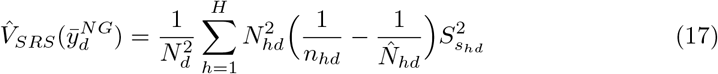

where 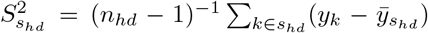 is the intersection sample variance and 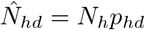. It is worthwhile to notice that (17) requires at least *n_hd_* ≥ 2 to allow 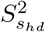 to be calculated.

The considerations regarding the sample size of the post-stratified estimator also apply to Eq. (11). That is, none of the intersection sample sizes *n_hd_* should be too small. Särndal et al (1992, Remark 10.7.2) suggests a minimum sample size of at least 10 observations in each intersection.

#### Relative precision of the nested group and π- estimators

Under stratified random sampling the approximate variance of the π-estimator shown in Eq. (2) takes the form

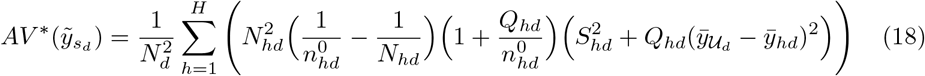

A proof is provided in the Appendix B.

The variance of the nested group estimator is

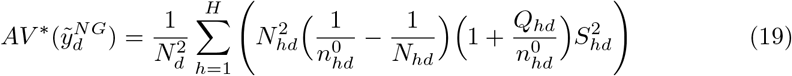

which implies that 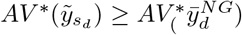 always. The smaller the domain and the larger the differences among the strata means, the larger the gain in precision due to the use of the nested group estimator is. The drawback is that the nested group estimator is more *demanding* in terms of sample size, as it requires to have at least one sample in each intersection 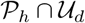.

The minimum of the *V*(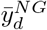) for a certain domain *𝒰_d_*, given a fixed total sample size *n*, is obtained when:

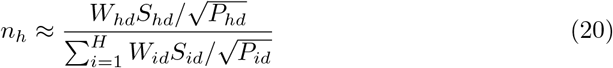

Eq. (20) corresponds to the approximate optimum allocation for the nested estimator for the the domain *𝒰_d_* (cf. Cochran 1977, Chap.5), and contrasts to the traditionally used Neyman optimum allocation for stratified random sampling (Neyman 1934):

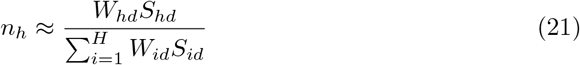

A proof on the development of Eq. (20) is provided in Appendix C.

## 5 Numerical tests of estimators

The properties of the above mentioned estimators were examined in a simulation study. We chose here a spatially explicit domain subject to stratified systematic sampling. Let us consider a population *𝒫* consisting of 320 non-overlapping square units each with area 1 ha labeled *k = 1,…,320*, so that *𝒫 = {u_1_,…, u_k_,…, u_320_ }*. The population *𝒫* is partitioned into 2 strata *a* and *b*, of size 150 and 170 ha, respectively. Let *Y* be the variable of interest. Values of *Y* have been randomly assigned, according to a normal distribution with mean and variance parameters as described in Table 2, to each population unit in the 2 strata. Further, let us assume that a random stratified sample s of size 35 has to be drawn without replacement from the population based on the two strata *a* and *b*. That is, a random sample sa of size na has to be selected from the stratum a and a random sample *s_b_* of size *n_b_* from the stratum *b*, so that *n_a_* + *n_b_* = 35. The population has also been partitioned into 2 domains, *A* and *B*, of size 96 and 224 ha, respectively (Fig. 1). Population statistics are displayed in Table 2.

**Fig. 1.**
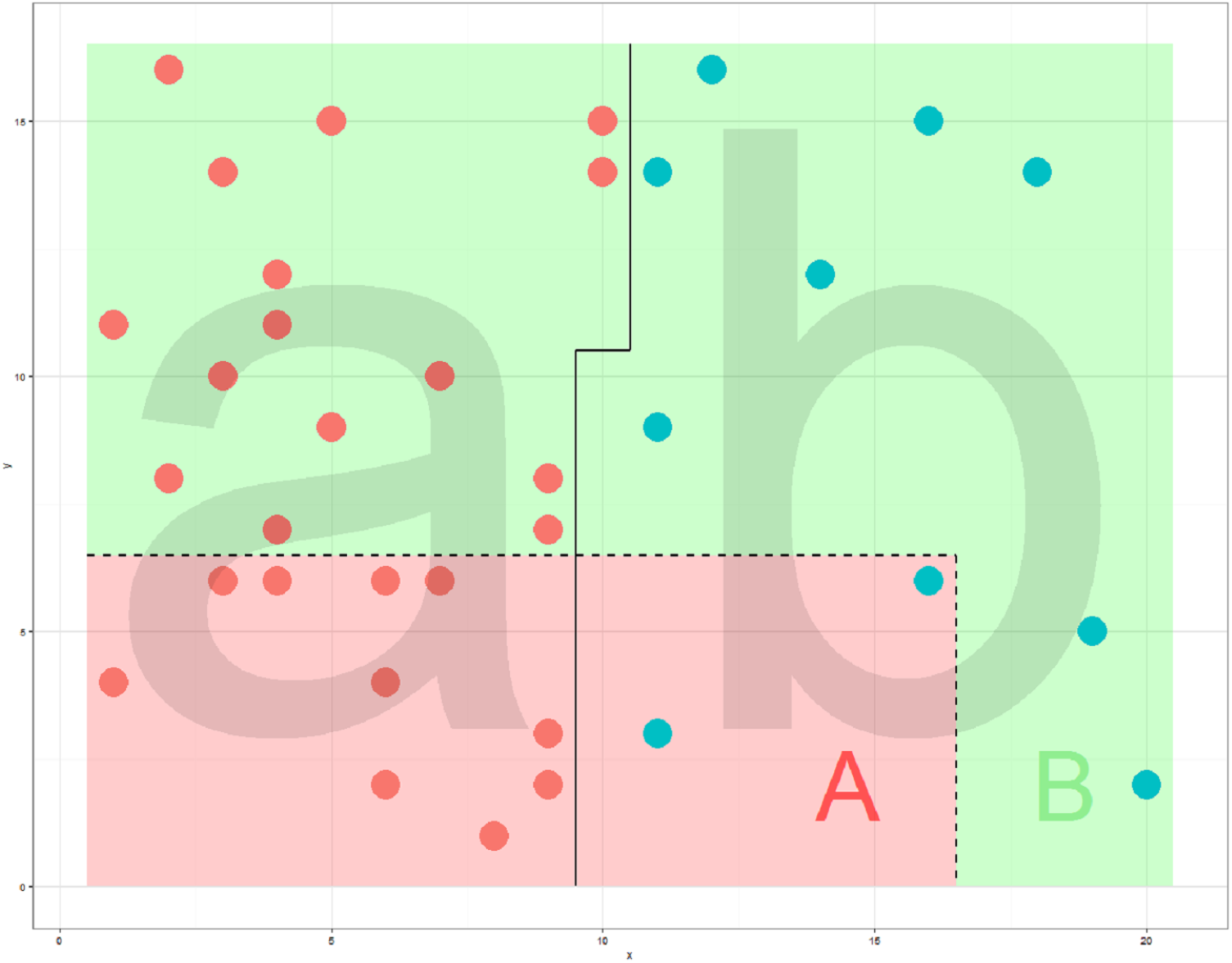
The area of study consists of a rectangular region of 320 ha, subdivided into 320 non-overlapping square areas of 1 hectare. Sampling- strata *a* and *b* are divided by the solid line, while domains *A*, in red and *B*, in green are divided by the dotted line. For illustration purposes, a simple random sample is shown here independently drawn from each sampling-stratum, with allocation not proportional to the size of the strata (in this a particular allocation *n_a_* = 25, red dots, and *n_b_* = 10, blue dots).

**Table 2.** Statistics of strata and domains used in the simulation study

| Population total<br>$N = 320$<br>$s = 35$ | |
| --- | --- |
| strata | domains |
| $N_a = 150$ | $N_A = 96$ |
| $N_b = 170$ | $N_B = 224$ |
| $\bar{Y}_a = 15.6$ | $\bar{Y}_A = 19$ |
| $\bar{Y}_b = 24$ | $\bar{Y}_B = 20.5$ |
| $S_a^2 = 26.6$ | $S_A^2 = 36.1$ |
| $S_b^2 = 13.9$ | $S_B^2 = 36.9$ |

The domain averages *Ȳ_A_* and *Ȳ_B_* were estimated from the sample *s*. The number of plots selected in strata *a* and *b* had not been initially defined and 16 different plot allocation strategies has been tested, with *n_a_* ϵ (10, 25) and *n_b_* = *n* − *n_a_*. For each allocation 10^5^ independent sample replicates of *n* = 35 were drawn and from each of those samples the domain means were estimated. Given the large number of samples and allocation regimes, we simulated parallel code with the package doParallel (Revolution Title Suppressed Due to Excessive Length Analytics and Weston 2015) in R (R Core Team 2016), using a cluster of 16 CPU and 30 GB of RAM provided by the FAO-hosted SEPAL platform (SEPAL 2016). We contrasted the following estimators:

1. the domain sample arithmetic mean, Eq. (4)
2. the π-estimator, Eq. (9)
3. the nested group estimator, Eq. (14)

The means of the 10^5^ sample replicates, estimates for the 2 domains for each plot allocation and for each estimator are presented in Fig. 2.

**Fig. 2.**
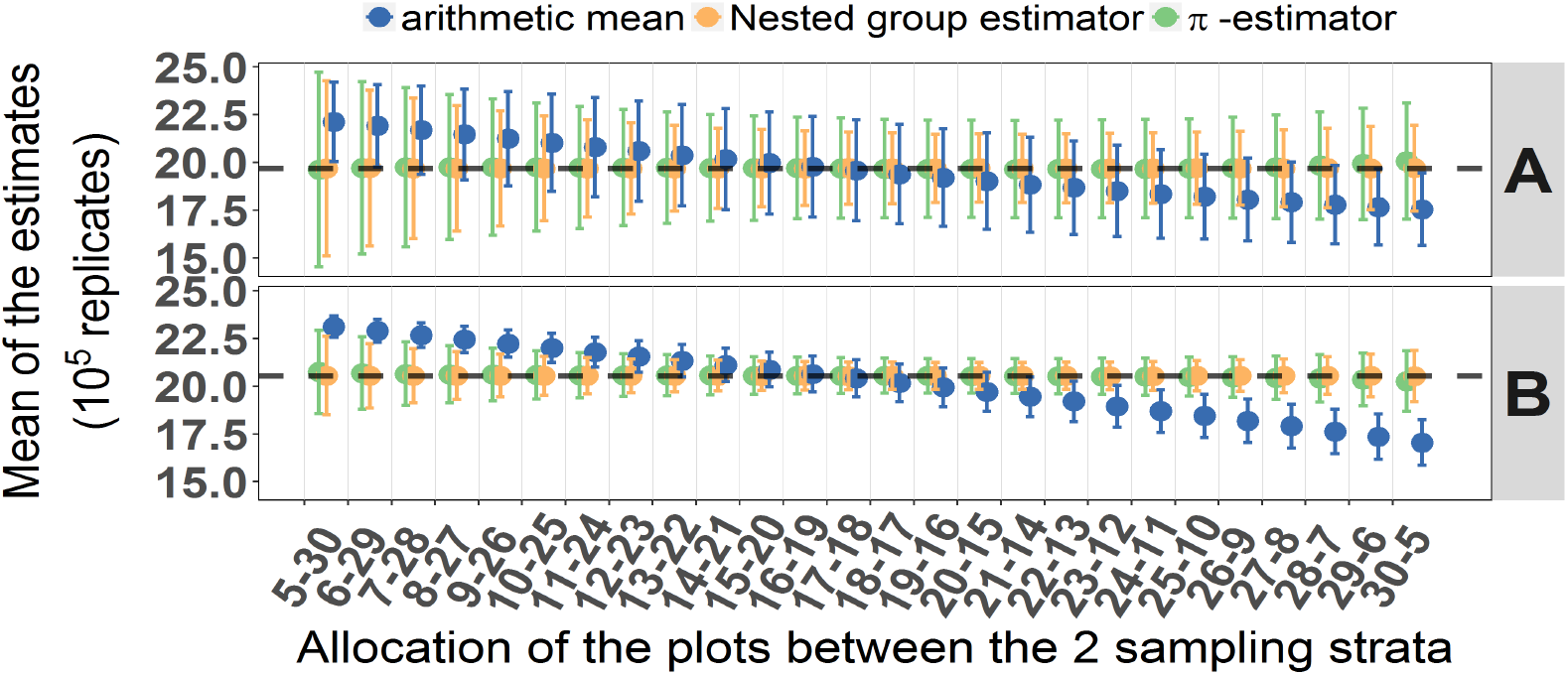
Means of the estimates for the domains *A* and *B*, by allocation and estimator. The grey dashed horizontal line is the real mean of the domains

As expected, the arithmetic sample mean proves biased, whenever the sample allocation is far from being proportional to the strata size (which corresponds to the allocation “16-19”). 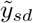 and 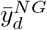 are approximately unbiased. The MSE is defined as the sum of the variance and the squared bias. In our example, the nested group estimate presents smaller MSE than the π-weighted estimate in all allocation scenarios (Fig. 3), and both always smaller than the MSE for the arithmetic mean. The numerically obtained optimum allocation strategy, would correspond to “22-13”(π-weighted) *vs.* “19-16” (nested group) when minimizing MSE for the “A” domain mean, and “21-14” for both estimators in the MSE minimization of the “B” domain.

**Fig. 3.**
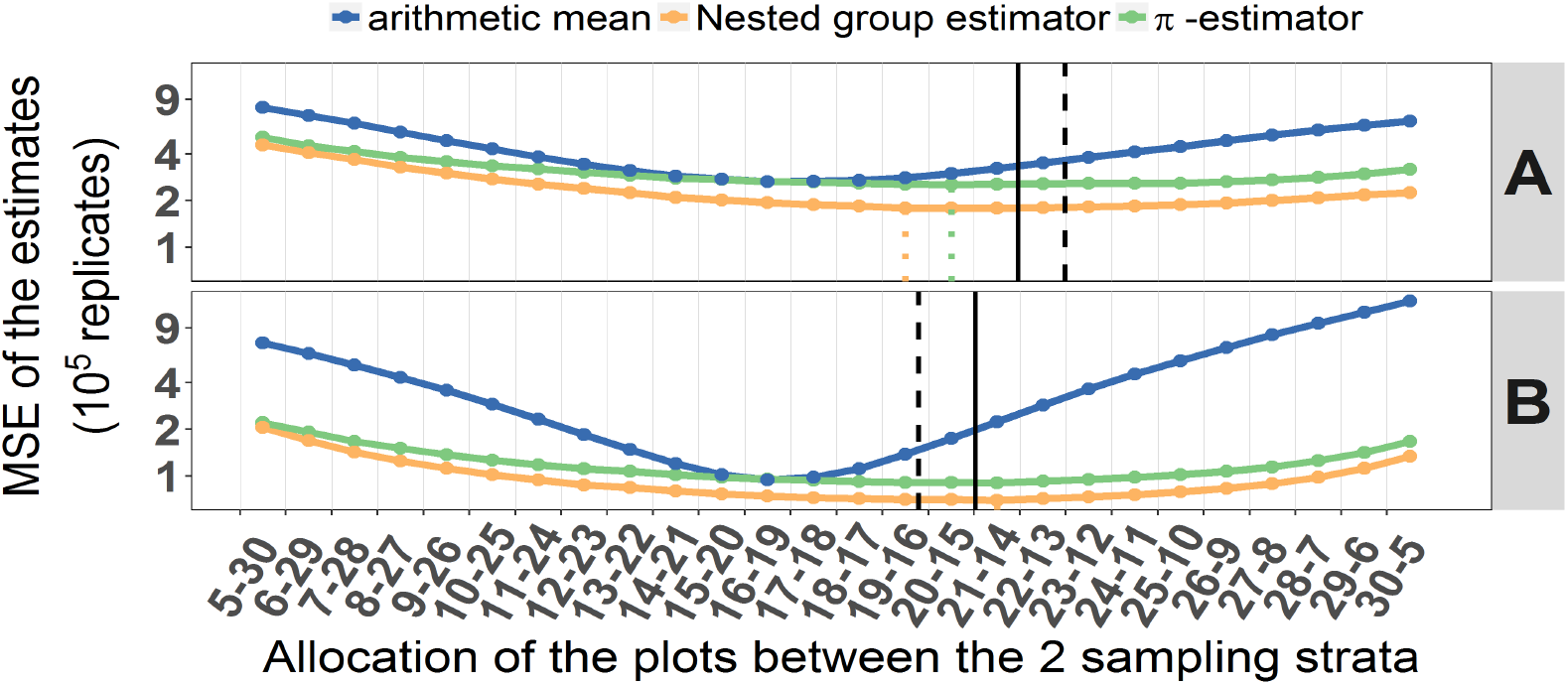
Mean squared errors of the estimates for the domains *A* and *B*, by allocation and estimator. Dotted green and orange vertical lines reflect the allocations at which the MSE of the domain mean is minimized in the π- *vs*. nested group estimators, respectively. Black solid and dashed vertical lines correspond to nested group (Eq. (20)) *vs*. Neyman (Eq. (21)) theoretical optimum allocations. The y-axis is log-transformed for better visualization.

Only by teasing apart the two components of the MSE, however, can one disentangle the role of bias and precision in each of the estimates. This becomes particularly relevant for REDD+ countries aiming to follow IPCC (2003) guidelines, recommending the minimization of uncertainty (i.e., precision) as much as possible. Precision (i.e., variance) estimates in the numerical tests show that *V*(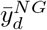) is always smaller than *V*(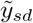) (Fig.4). However, the arithmetic mean presents smaller variance values in those allocation strategies far from the optimum. Minimum variances for both π-weighted and nested group estimators match the MSE values and indicate that for these two estimators the vast majority of the MSE is due to variance (and hence, reinforcing the unbiasedness of these two estimators).

**Fig. 4.**
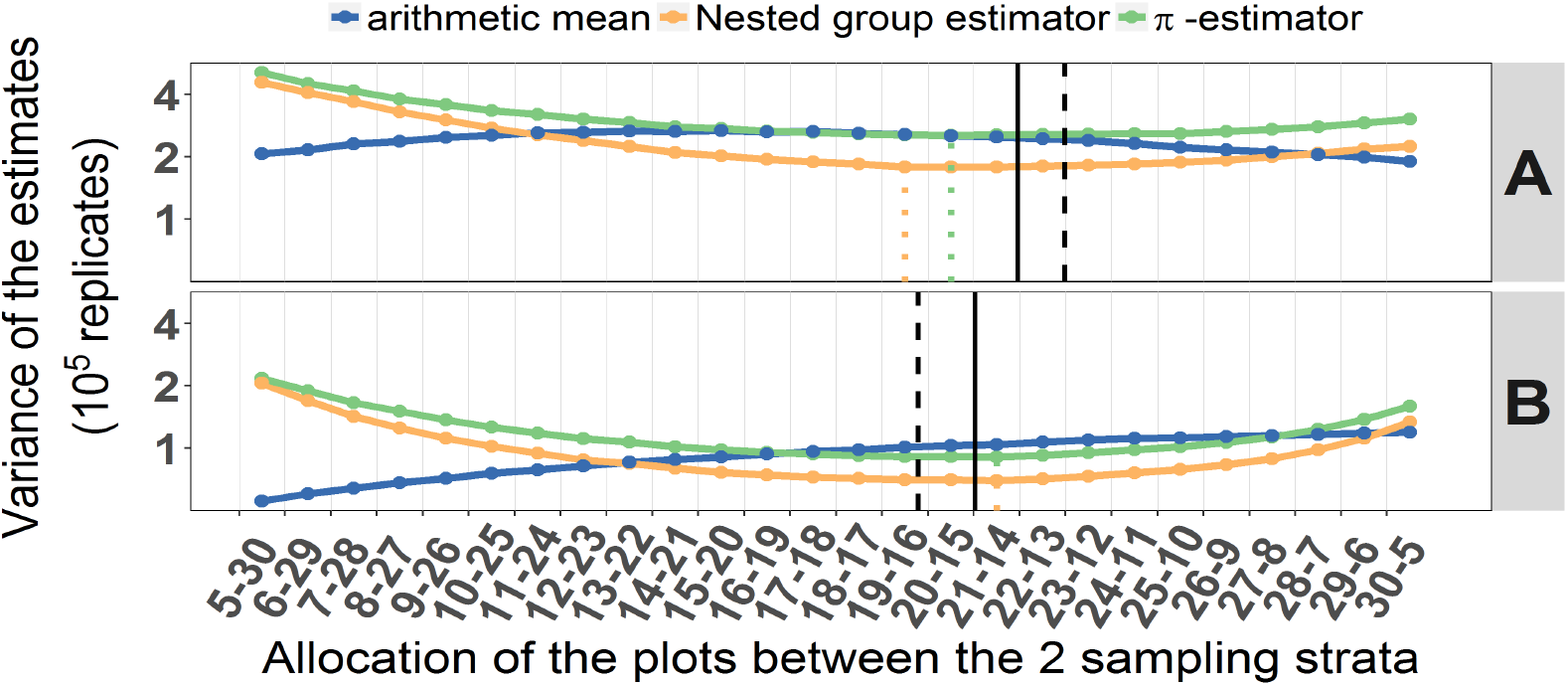
Variance of the estimates for the domains *A* and *B*, by allocation and estimator. Dotted green and orange vertical lines reflect the allocations at which the variance of the domain mean is minimized in the π- *vs*. nested group estimators, respectively. Black solid and dashed vertical lines as in Fig. 3. The y-axis is log-transformed for better visualization.

## 6 Discussion

The amount of information which large-scale environmental surveys (such as national forest inventories) are expected to produce has considerably increased over time (McDonald 2003). In the case of GHG emissions reporting, this expectation might include the production of estimates for a certain number of domains of interest (such as land use categories) (Tubiello et al 2014), often identified after and independently of the forest inventory (IPCC 2003). But obtaining accurate and unbiased estimates for such domains is often a non trivial exercise, especially in the presence of complex survey designs (Lehtonen and Pahkinen 2004).

Thus, when the domains are spatially explicit (i.e. each population unit is classified into a domain) and all the forest inventory sample units geo-referenced, it is always possible to assign automatically each of n sample units into a domain *𝒰_d_*, even when the information regarding the domains was not collected during the NFI. Often, however, the domains might not be spatially explicit and their sizes might have been estimated independently of the forest inventory, such as, for example, through an external, ancillary sample-based land use survey with a different sampling design. This can occur frequently during the preparation of a GHGI report, where activity data (areas of land use or land use changes) and emission factors (i.e., biomass, volume,‥) are typically compiled from disparate sources. This method for the representation of land in tabular form falls within the IPCC definitions of *approach 1* and *approach 2,* as described in IPCC (2003, Chap. 2) and IPCC (2006, Vol. 4, Chap. 3), and it is widely used for the estimation of land use and land use changes. An example of this is the case in which the absolute sizes of land use categories are independently estimated through visual (or augmented visual) interpretation of a sample of remote imagery (Bey et al 2016).

In this case, in order to estimate the domain averages it is first of all necessary to classify each of the n forest inventory sample units into one of the spatially-implicit domains *𝒰_d_*. To prevent any possible bias, re-classification needs to be done using the same methodology adopted to estimate the domain sizes. If, for example, these have been estimated through sample-based visual interpretation, then the *n* sample units need to be interpreted using the same type of imagery and, possibly, the same interpreters of the land use survey. The domain mean estimator to be used is the estimator 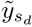 in Eq. (1), which under a simple random or systematic sampling corresponds to the domain sample mean Eq. (4) and under a stratified random or stratified systematic sampling takes the form of Eq. (9).

In an attempt to offer a set of rules for the selection of adequate domain estimators, we propose a decision tree (Fig. 5) that aims to guide forestry academics and NFI assessment teams through each step depending on:

1. the stratified nature of the NFI design
2. the ancillary nature of the domain sizes
3. whether the domains are spatially implicit or explicit
4. whether the sampling (per stratum) is SRS or systematic

**Fig. 5.**
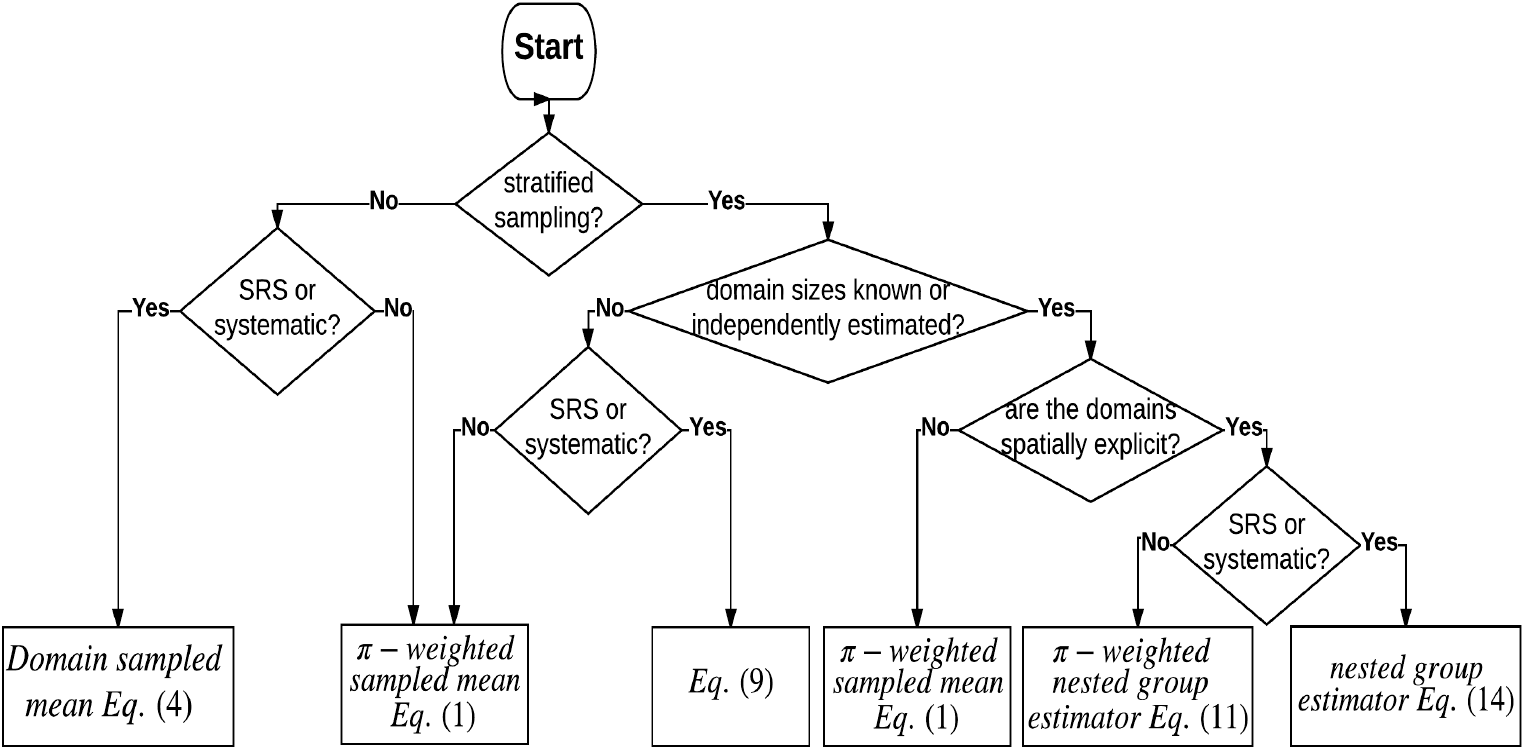
Decision tree for the selection of the adequate domain mean estimator

Overall, π-weighted estimators should be considered whenever the sampling per stratum (∀*H* ≥ 1) does not follow SRS or systematic designs, or else when the domains are spatially implicit and independently estimated. Under a SRS or systematic design, when *H* = 1 or *H* > 1 the domain sample mean Eq. (4) or Eq. (9) are recommended, respectively. Finally, if *H* > 1, domain sizes are independent from the NFI and spatially explicit, nested group estimators outperform the others, using the π-weighted version (Eq. (11)) if the per stratum samples were not SRS or systematically taken.

The use of nested estimators, thus, should ideally be known in order to facilitate allocation strategies previous to the design of the forest inventory. Notice that Eq. (20) keeps a striking similarity to the optimal allocation equation of Cochran (1977, p. 98) (Schreuder et al 1993). But instead, variables per stratum *h* are accounted on a per stratum/domain intersection *hd*, and Cochran’s costs of sampling a unit in stratum h are replaced by *P_hd_*. Hence, when the stratum is filled by the domain, *P_hd_ = 1* and optimum allocation corresponds to Neyman’s (21). Otherwise, *P_hd_* < 1 will effectively rescale the weight assigned to individual sampling units in the s tratum, akin to a reduction in cost per sampling unit.

In the simulation exercise, Eq. (20) outperforms Neyman’s Eq. (21) for both domains as references. However, it is still an approximation, as shown in Appendix C, largely driven by the limitations imposed in the exercise, when *n_h_* is small. Although the situation under which the application of nested group estimators and the nested group optimal allocation strategy might seem rare, many countries find themselves in practice conducting international reporting where per ha. emission factors are calculated from stratified inventories, and the calculation of domain sizes comes from independent, spatially explicit sources. As shown above, this is where nested group estimators may provide opportunities to reduce both bias and sampling error, outperforming not only the often misused domain arithmetic means, but also the classical unbiased π-weighted estimators.

While the nested group approach might seem the ideal one given its performance in our example, it is subject to several assumptions that often may not be as realistic. First, *n_hd_* ≥ 2 is a necessary condition to obtain a measure of precision; second, and most important, domain sizes are assumed to be error-free. This assumption, although often taken, is far from realistic, since spatially explicit obtained information always contains errors, even if only sampling ones. Despite this last assumption, most international reporting so far from countries to the UNFCCC has been known to observe it, at least for static pictures (i.e., area changes are calculated with their errors). In such a case, the use of nested estimators might be the best option for REDD+ reporting.

## Footnotes

1 From IPCC (2003): “Estimates should be accurate in the sense that they are systematically neither over nor under true emissions or removals, so far as can be judged, and that uncertainties are reduced so far as is practicable”

## Acknowledgements

The authors want to thank specially Becky Tavani and Julian Fox from the Forestry Department in FAO, and the whole UN-REDD program for their insights and continuous support in the preparation of this document.

## Appendices

## A The bias of the domain sample mean under a random stratified sampling

Similarly to §4, let us consider the case in which a stratified random sampling is carried out on the population *𝒫*, based on the *H* sampling-strata. Let denote *s_hd_* the intersection of the domain *𝒰_d_* and the sample drawn in the sampling-stratum *𝒫_h_*, and *n_hd_* the size of this sub-sample. The mean of *Y* for *s_d_* can be written as:

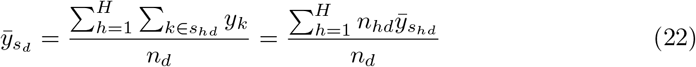

where 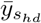 is the sample mean of the sample drawn in the in the stratum *𝒫_h_*, and belonging to the domain *𝒰_d_*. The number *n_hd_* of sample units falling into 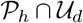 varies depending on the sampling strategy and on the sample allocation adopted. If a simple random or systematic sample is taken in each sampling-stratum the expected value of the number of units *n_hd_* that are in *s_d_* is given by:

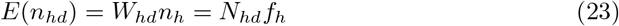

where *W_hd_* = *N_hd_/N_h_* is the relative size of the intersection 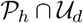 within the stratum *𝒫_h_*. The expected value of the number of sample units *n_d_* that happen to fall into the domain *𝒰_d_* can be expressed as a linear function of the expected number of sample units in each intersection:

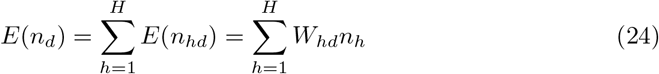

Denote by *A_d_* the event *E*(*n*_*d*_) ≥ 1. If n is large enough *Pr*(*A*_*d*_) is close to 1. Under the condition *A_d_*, the expected value of 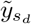 for the domain *𝒰_d_* is given by

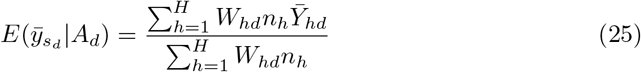

We will now assume that *A_d_* will certainly occur. Since the population mean of the domain *𝒰_d_* can be written as:

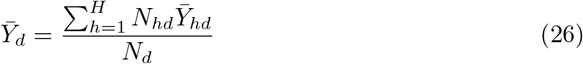

the bias of the arithmetic mean 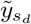 for the domain *𝒰_d_* therefore amounts to:

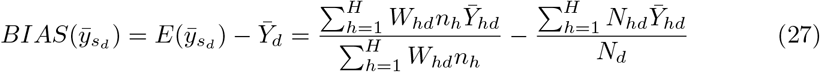

Under a stratified random sampling with a sample allocation proportional to the size of the strata, where *n_h_ = nN_h_/N = N_h_ƒ*, the arithmetic mean proves unbiased since

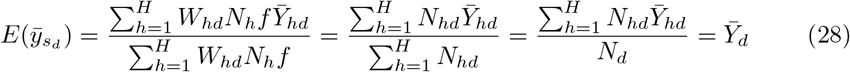

## B A better approximation of the variance of the π-estimator under a stratified random sampling

Under a random stratified sampling with *H* strata, indexed with the letter *h*, (where *h = 1,…, H)* (2) can be re-expressed as

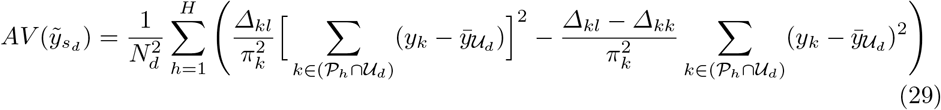

and

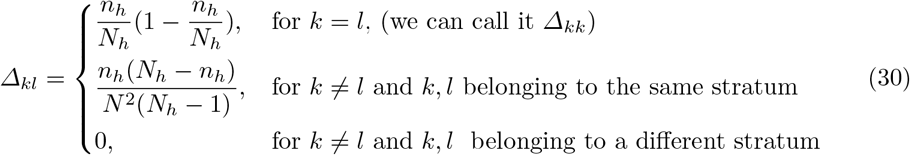

therefore

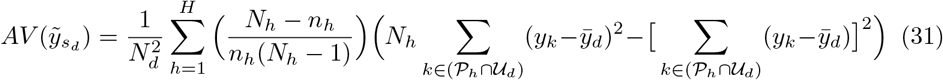

decomposing the variance we obtain:

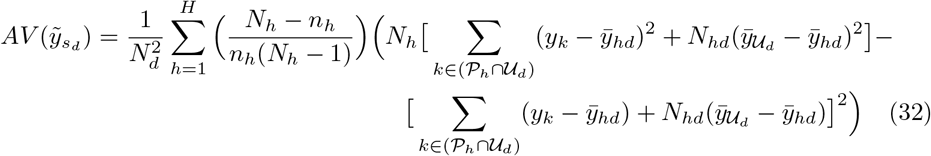

hence

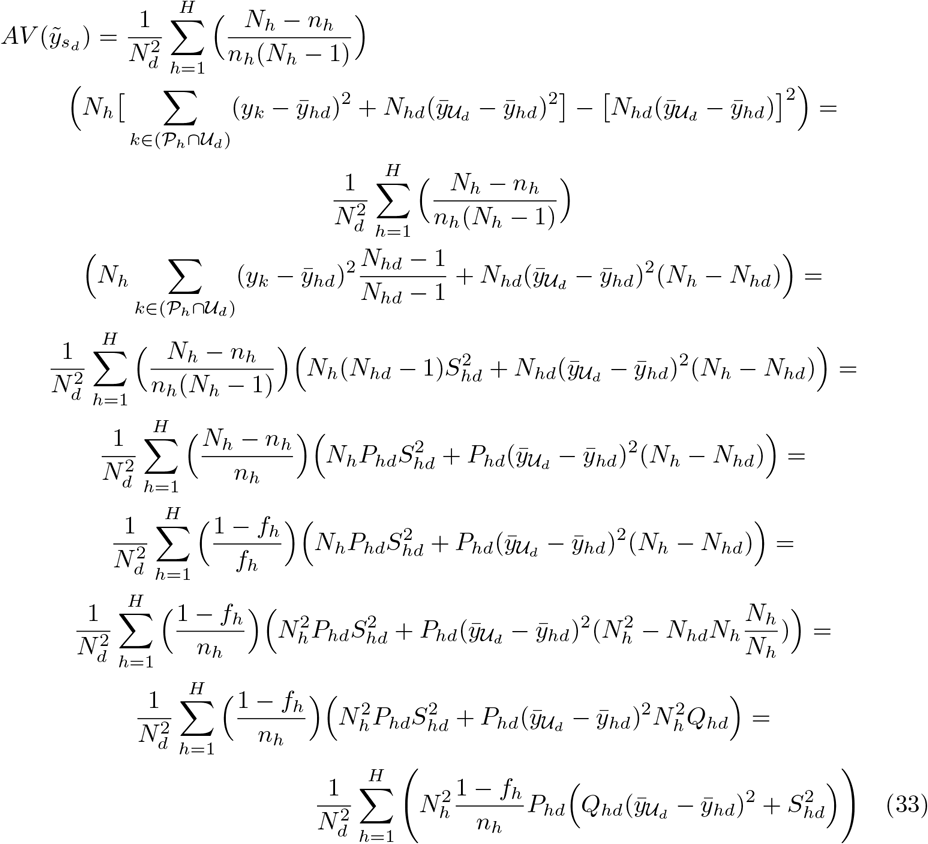

where 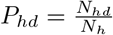, *Q_hd_ = 1 − P_hd_* and *ƒ_h_ = n_h_/N_h_*.

By analogy with Särndal 10.3.15, (33) can also be expressed as:

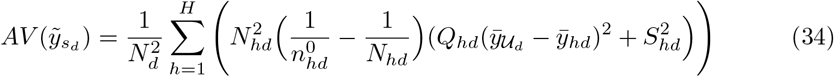

where 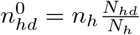.

By conditioning on the domain sample size, similarly to the development in §10.4, of Särndal et al (1992), we get a closer approximation to *V*(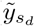)

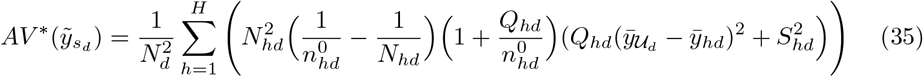

## C Optimum sample allocation for domain estimation under random stratified sampling

*Theorem:* The sample sizes *n_1_,…, n_h_,…, n_H_* that minimize the 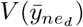 under a random stratified sampling, subject to the constraint *n_1_ +… + n_h_ +… + n_H_ = n* are approximated by

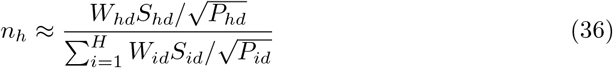

*Proof* Under a random stratified sampling, neglecting the finite population correction *1/N_hd_*, (19) can be written as

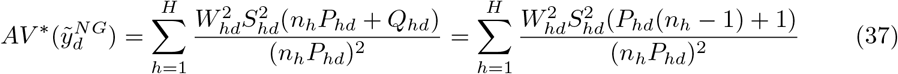

Where *Q_hd_* = 1 - *P_hd_* and *W_hd_* = *N_hd_/N_d_*. Under the condition *n_h_* >> 0, *n_h_* - 1 ≈ *n_h_* and (*n_h_P_hd_* + 1)/*n*_*h*_*P*_*hd*_ ≈ 1. (37) therefore simplifies to

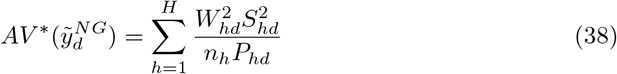

The minimum of this function can be found using the method of Lagrange multipliers. The Lagrange function is:

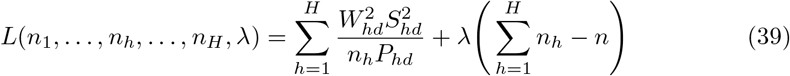

For *h* = 1,…, *h*,…, *H*, where λ is the Lagrange multiplier. The partial derivatives of this function are

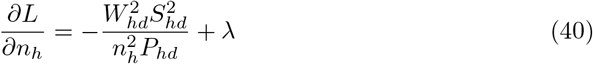

By setting the partial derivatives of this function equal to zero we obtain

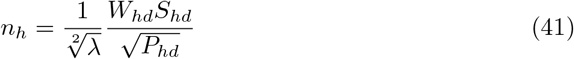

By summing these equations over *h*:

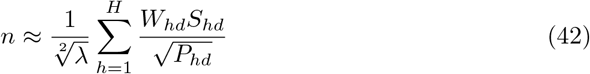

Thus,

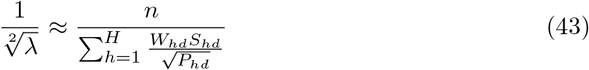

and

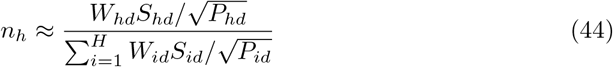

## References

Bechtold WA, Patterson PL (2005) The Enhanced Forest Inventory and Analysis Program - National Sampling Design and Estimation Procedures. USDA Forest Service General Technical Report SRS-80:85

Bey A, Sánchez-Paus Díaz A, Maniatis D, Marchi G, Mollicone D, Ricci S, Bastin JF,Moore R, Federici S, Rezende M, Patriarca C, Turia R, Gamoga G, Abe H, Kaidong E,Miceli G (2016) Collect Earth: land use and land cover assessment through augmented visual interpretation. Remote Sens-Basel 8(10):807, DOI 10.3390/rs8100807

Chanbers RL (2011) Which sample survey strategy? a review of three different approaches. Pak J Stat 27(4):337–357

Cochran WG (1977) Sampling Techniques 3rd ed. Wiley publication in applied statistics,John Wiley&Sons: New York

Gregoire TG, Valentine HT (2008) Sampling Strategies for Natural Resources and the Environment. Chapman&Hall/CRC

Holt D, Smith TF (1979) Post stratification. J R Stat Soc Ser A-G 142(1):33–46, DOI 10.2307/2344652

IPCC (2003) Good Practice Guidance for Land Use, Land-Use Change and Forestry. Institute for Global Environmental Strategies, Kanagawa

IPCC (2006) Guidelines for National Greenhouse Gas Inventories. Volume 4. Agriculture, Forestry and Other Land Use. Institute for Global Environmental Strategies, Kanagawa

Kish L (1980) Design and estimation for domains. The Statistician 29(4):209–222, DOI 10.2307/2987728

Köhl M, Magnussen SS, Marchetti M (2006) Sampling Methods, Remote Sensing and GIS Multiresource Forest Inventory. Tropical Forestry, Springer Berlin Heidelberg,DOI 10.1007/978-3-540-32572-7

Lehtonen R, Pahkinen E (2004) Practical methods for design and analysis of complex surveys. John Wiley&Sons, DOI 10.1002/0470091649

Lohr SL (2009) Sampling: Design and Analysis. Advanced (Cengage Learning),Brooks/Cole Pub. Co.: Boston

Mandallaz D (2008) Sampling Techniques for Forest Inventories. Chapman&Hall/CRC Applied Environmental Statistics, Taylor&Francis, DOI 10.1201/9781584889779

Maniatis D, Mollicone D (2010) Options for sampling and stratification for national forest inventories to implement REDD’ under the UNFCCC. Carbon Balance Manag 5(1):9, DOI 10.1186/1750-0680-5-9

McDonald TL (2003) Review of environmental monitoring methods: Survey designs.Environmental Monitoring and Assessment 85(3):277–292, DOI 10.1023/A:1023954311636

Neyman J (1934) On the two different aspects of the representative method: the method of stratified sampling and the method of purposive selection. J R Stat Soc 97(4):558–625, DOI 10.2307/2342192

Pacificador Jr AY (1997) The sample mean under stratified random sampling. Philipp Stat 46:73-82

R Core Team (2016) R: A Language and Environment for Statistical Computing. R Foundation for Statistical Computing, Vienna, Austria, URL https://www.R-project.org/

Rao JN, Molina I (2015) Small Area Estimation. John Wiley&Sons, DOI 10.1002/9781118735855

Revolution Analytics, Weston S (2015) doParallel: Foreach Parallel Adaptor for the ‘parallel’ Package. URL https://CRAN.R-project.org/package=doParallel, r package version 1.0.10

Särndal CE, Lundström S (2005) Estimation in Surveys with Nonresponse. John Wiley&Sons, DOI 10.1002/0470011351

Särndal CE, Swensson B, Wretman J (1992) Model Assisted Survey Sampling. Springer-Verlag, DOI 10.1007/978-1-4612-4378-6

Schreuder H, Gregoire T, Wood G (1993) Sampling Methods for Multiresource Forest Inventory. Wiley

Schreuder HT, Ernst R, Ramirez-Maldonado H (2004) Statistical techniques for sampling and monitoring natural resources. USDA Forest Service General Technical Report RMRS-GTR-126

Schulz B, Bechtold W, Zarnoch S (2009) Sampling and estimation procedures for the vegetation diversity and structure indicator. General technical report PNW, U.S. Dept.of Agriculture, Forest Service, Pacific Northwest Research Station, Portland, OR

SEPAL (2016) System for Earth Observation Data Access, Processing and Analysis for Land Monitoring. URL https://sepal.io, accessed: 2017-04-14

Thompson SK (2012) Sampling. John Wiley & Sons, Hoboken, NJ 10, DOI 10.1002/9781118162934

Tubiello F, Salvatore M, Cóndor Golec R, Ferrara A, Rossi S, Biancalani R, Federici S,Jacobs H, Flammini A (2014) Agriculture, forestry and other land use emissions by sources and removals by sinks. FAO Statistics Division, Food and Agriculture Organization, Rome ESS/14-02:89

